## Supplemental Table S1 and Figs. S1 and S2 for "Cancer malignancy is correlated with up-regulation of PCYT2-mediated glycerol phosphate modification of *α*-dystroglycan"

[1] Exploratory Research Center on Life and Living Systems (ExCELLS) and [2]Institute for Molecular Science (IMS), National Institutes of Natural Sciences, Okazaki, Japan
[3] Department of Structural Biology and Biomolecular Engineering, Graduate School of Pharmaceutical Scienc-es and [4]Department of Gastroenterology and Metabolism, Graduate School of Medical Sciences, Nagoya City University, Nagoya, Japan
[5] Center for Joint Research Facilities Support, Research Promotion and Support Headquarters, Fujita Health University, Toyoake, Aich, Japan
[6] Institute for Glyco-core Research (iGCORE), Gifu University, Gifu, Japan
[7] Laboratory of Microbiology and Immunology, Graduate School of Pharmaceutical Sciences, Chiba University, Chiba, Japan
8 Institute of Biological Chemistry, Academia Sinica, Taipei, Taiwan

**Table S1: Pathological characteristics and DG2 staining of the tissue microarray.**

| Position | Sex | Age | Anatomic Site | Pathology | Grade | Stage | DG2 |
|---|---|---|---|---|---|---|---|
| A01 | M | 65 | Bladder, urinary | Normal | null | null | − |
| A02 | F | 39 | Bladder, urinary | Transitional cell carcinoma | III | T2N0M0 | + |
| A03 | M | 61 | Bladder, urinary | Transitional cell carcinoma | III | T2N0M0 | + |
| A04 | F | 34 | Breast | Normal breast | null | null | + |
| A05 | F | 43 | Breast | Invasive ductal carcinoma | II | T3N1M0 | + |
| A06 | F | 75 | Breast | Invasive ductal carcinoma | III | T3N0M0 | + |
| A07 | M | 32 | Stomach | Normal stomach | null | null | + |
| A08 | M | 66 | Stomach | Adenocarcinoma | II | T3N0M0 | + |
| B01 | M | 47 | Stomach | Adenocarcinoma | III | T3N2M0 | + |
| B02 | F | 43 | Intestine, colon | Normal colon | null | null | − |
| B03 | M | 36 | Intestine, colon | Adenocarcinoma | III | T3N1M0 | + |
| B04 | M | 55 | Intestine, rectum | Adenocarcinoma | I~II | T3N1M0 | + |
| B05 | M | 30 | Kidney | Normal kidney cortex | null | null | + |
| B06 | F | 79 | Kidney | Cear cell carcinoma | null | T1N0M0 | + |
| B07 | M | 72 | Kidney | Cear cell carcinoma | null | T1N0M0 | + |
| B08 | M | 35 | Liver | Normal | null | null | + |
| C01 | M | 59 | Liver | Hepatocellular carcinoma | II | T3N0M0 | + |
| C02 | M | 19 | Liver | Hepatocellular carcinoma | I~II | T2N0M0 | + |
| C03 | M | 59 | Lung | Normal | null | null | + |
| C04 | M | 58 | Lung | Squamous cell carcinoma | II~III | T2N0M0 | + |
| C05 | F | 40 | Lung | Adenocarcinoma | II~III | T2N0M0 | + |
| C08 | M | 42 | Lymph node, axillary | Lymphoma, non-Hodgkin B cell lymphoma | null | null | + |
| D01 | F | 23 | Ovary | Normal | null | null | − |
| D02 | F | 43 | Ovary | Mucinous cystadenocarcinoma | III | T2N0M0 | + |
| D03 | F | 58 | Ovary | Endometrioid adenocarcinoma | III | T2N0M0 | + |
| D04 | F | 58 | Pancreas | Normal | null | null | + |
| D05 | F | 49 | Pancreas | Adenocarcinoma | II | T3N1M1 | + |
| D07 | M | 73 | Prostate | Normal | null | null | + |
| D08 | M | 63 | Prostate | Adenocarcinoma | III | T3N0M0 | + |
| E01 | M | 60 | Prostate | Adenocarcinoma | II | T2N0M0 | + |
| E03 | M | 34 | Penis | Squamous cell carcinoma | I | T1N0M0 | + |

| | | | | | | | |
|---|---|---|---|---|---|---|---|
| E04 | M | 73 | Skin | Melanoma | null | null | + |
| E05 | F | 46 | Uterus | Normal | null | null | − |
| E06 | F | 41 | Uterus, cervix | Squamous cell carcinoma | I~II | T1N1M0 | + |
| E07 | F | 53 | Uterus, endometrium | Adenocarcinoma | II~III | T1N1M0 | + |

Staining with DG2 was classified as negative; − and positive; +.

### Figure S1

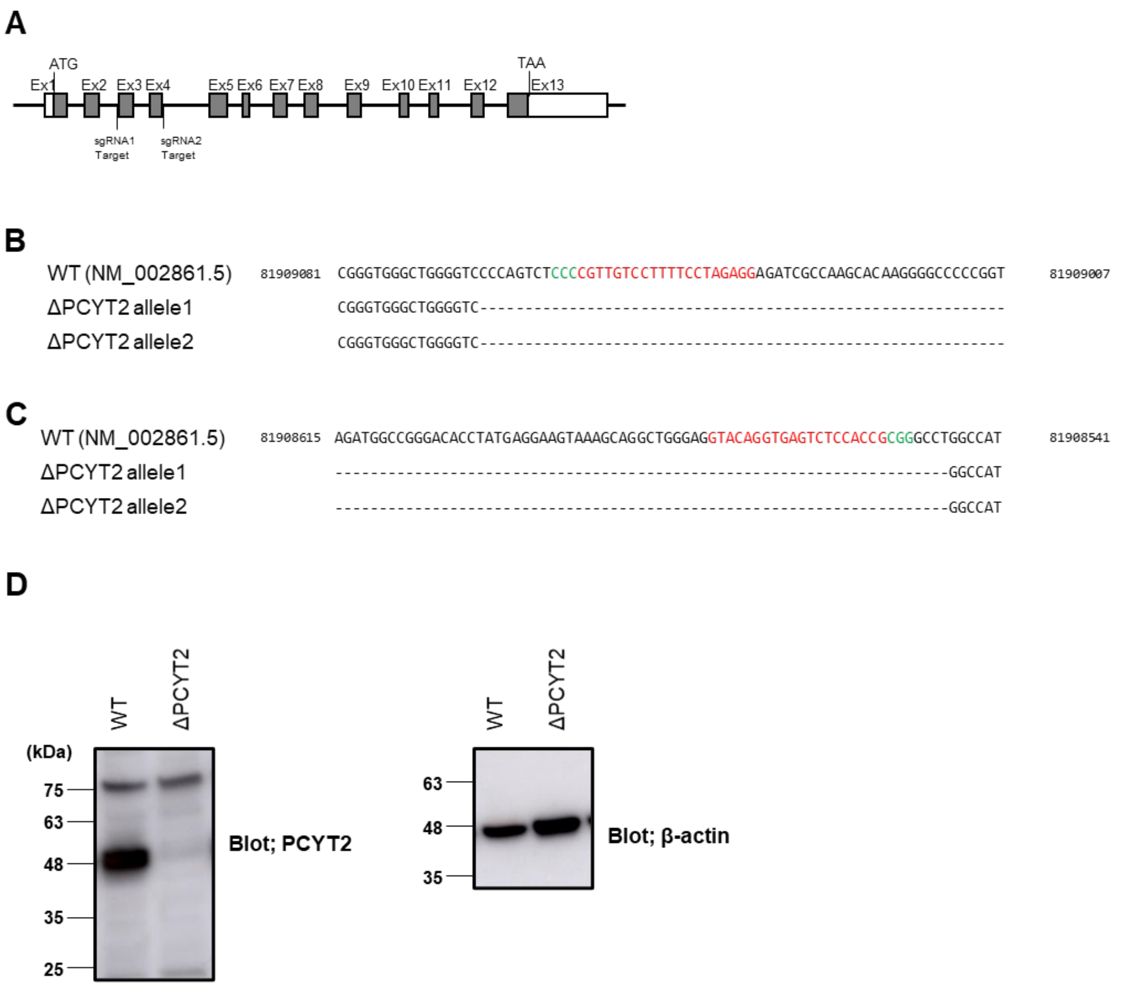


**Figure S1: Editing of PCYT2 gene by the CRISPR/Cas9 system.** (A) Schematic representation of the genomic target sites in the PCYT2 gene. Exons, indicated by rectangles, are numbered from Ex1 to Ex13. The coding portions of the gene are shaded. The open portions of exons 1 and 13 represent the 5′ UTR and 3′ UTR, respectively. (B, C) Genotyping results of PCYT2-KO cells. Genomic DNAs from wild-type (WT) and KO cells were isolated, and PCR amplicons flanking the CRISPR/Cas9-targeted regions were sequenced. Sequence alignments of bases (B) 81909081-81909007 and (C) 81908615-81908541 of the PCYT2 gene locations are shown. The sequences of gRNAs and PAM are labeled in red and green; -, nucleotide deletion. Numbering is based on the entire PCYT2 gene sequence using the reference gene RefSeq NM_002861.5. (D) PCYT2 protein expression level in WT and PCYT2-KO (ΔPCYT2) cells were analyzed by immunoblotting with anti-PCYT2 and anti-β-actin antibodies. The equal amount of protein in each sample was adjusted and loaded for Western blotting.

**Figure S2**

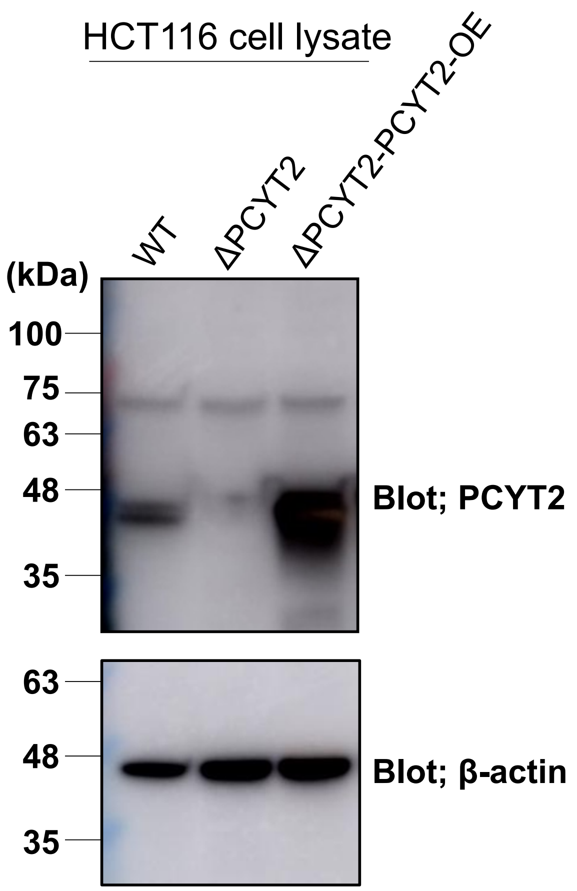


**Figure S2: PCYT2 expression in HCT116 wild-type, PCYT2-KO, and PCYT2-rescue cells.** Cell lysates containing equal amounts of total proteins that were prepared from HCT116 wild-type (WT), PCYT2-KO (ΔPCYT2), and PCYT2-KO rescue (ΔPCYT2-PCYT2-OE) cells were subjected to immunoblot analysis using anti-PCYT2 and anti-β-actin antibodies.
